# Detecting modifications in proteomics experiments with Param-Medic

**DOI:** 10.1101/496349

**Authors:** Damon H. May, Kaipo Tamura, William S. Noble

## Abstract

Searching tandem mass spectra against a peptide database requires accurate knowledge of various experimental parameters, including machine settings and details of the sample preparation protocol. In some cases, such as in re-analysis of public data sets, this experimental metadata may be missing or inaccurate. We describe a method for automatically inferring the presence of various types of modifications, including stable-isotope and isobaric labeling, tandem mass tags, and phosphorylation, directly from a given set of mass spectra. We demonstrate the sensitivity and specificity of the proposed approach, and we provide open source Python and C++ implementations in a new version of the software tool Param-Medic.

## 1 Introduction

Properly searching shotgun proteomics data requires accurate metadata. For example, searching with sub-optimal tolerances for precursor and fragment mass error can lead to a dramatic loss of sensitivity at a given false discovery rate.^1^ Search parameters must also account for any appropriate modifications, such as phosphorylation or isotopic labeling. If such modifications are not included in the search, then modified peptides cannot be identified. On the other hand, if variable modifications are included in the search parameters but are not present in the sample, then the search space is unnecessarily expanded, and search sensitivity at a given false discovery rate threshold suffers.

In most cases, the appropriate modification-related parameters for a particular mass spectrometry run are known to the person performing the search. When a lab is searching data that it produced, presumably information is easily obtained about which runs were enriched for phosphorylated peptides or included labels for quantitation and which runs did not. However, in other situations this information may not be readily available. For example, when searching data downloaded from a public repository, the appropriate metadata may not be available, may not be trustworthy, or may not apply to all of the runs in an experiment. Repositories performing re-searches of submitted data may also lack this information. Also, a researcher may want to perform a quick assessment of their own data in order to determine whether an expected modification (e.g., phospho-enrichment) is present and detectable.

We previously introduced Param-Medic, ^1^ a software tool that enables automatic inference of precursor and fragment mass accuracy parameters. In this work, we expand Param-Medic's functionality to automatically detect the presence of the following modifications: enrichment for phosphorylated peptides, SILAC labeling,^2^ and two different isobaric labeling schemes (iTRAQ^3,4^ and TMT^5,6^). We demonstrate Param-Medic’s sensitivity and specificity on each of these tasks by using publicly available data. We make this functionality available in both open source implementations of Param-Medic: the standalone Python tool (http://noble.gs.washington.edu/proj/param-medic) and the implementation integrated into the Crux proteomics toolkit (http://crux.ms).

## 2 Methods

For each of three general types of modifications—phosphorylation, isobaric labels, and SILAC labels—we developed a statistic that aims to distinguish between runs that contain that modification and runs that do not. Param-Medic computes each of these statistics and applies an empirically determined threshold to each one to determine the presence or absence of each type of modification. We developed and evaluated each classifier using public data from the MassIVE repository.^7^ Below, we describe the statistics, the data sets, and the methods for setting decision thresholds and evaluating performance.

### 2.1 Detecting Phospho-enrichment

Studying phosphorylation in the proteome requires enrichment of phosphorylated proteins.^8^ In phospho-enriched runs, phosphorylated peptides often contain a peak representing a neutral loss of H_3_PO_4_,^9^ occurring at an *m/z* value corresponding to a singly-charged mass ~98 Da less than the precursor mass.

Accordingly, Param-Medic assesses whether a given run is phospho-enriched by quantifying the observed signal corresponding to a loss of H_3_PO_4_. Specifically, Param-Medic assesses the proportion of total fragment signal accounted for by fragments falling in a nominal mass bin (with width 1.000507 Da, corresponding to the distance between the centers of two adjacent peptide mass clusters^10^) representing a 98-Da loss. This proportion is converted to a Z-score by comparing with the proportions of signal in eight nearby control bins (nominal *m/z* differences from precursor: −20, −15, −12, −10, 10, 12, 15, 20). If the Z-score exceeds an empirically determined threshold, then the run is determined to be phospho-enriched, and Param-Medic suggests that searches performed on the run use an appropriate variable modification of 79.966331 Da on serine, threonine and tyrosine residues.

### 2.2 Detecting Isobaric Labels

Isobaric labeling systems have become very popular for relative and absolute quantitation of peptides. Param-Medic now supports automatic detection of two labeling schemes: iTRAQ and TMT. Both iTRAQ and TMT use isobaric labels, with the intensity of reporter ions of fixed *m/z* ratio providing a measure of abundance in each sample.

To determine whether isobaric labels of a particular type (whether iTRAQ or TMT) are present, Param-Medic assesses the proportion of total MS/MS fragment signal that is accounted for by a list of nominal reporter ion *m/z* values for each type of label, as well as for a set of control *m/z* values (111, 112, 120, 122, 123, 124, 125, 133, and 134 *m/z*). Then, separately for iTRAQ and for TMT, Param-Medic calculates a t-statistic describing the extent to which the signal from all reporter ions for the label type is higher than the signal from the controls in order to determine whether the label responsible for the reporter ions is present. Finally, Param-Medic compares the *t*-statistic to an empirically determined threshold of 1.5 for iTRAQ labeling and 2.0 for TMT labeling.

Each labeling method comes in multiple varieties. For example, iTRAQ has both 4-plex and 8-plex versions, where the 4-plex reporter ions are a subset of the 8-plex reporter ions. Accordingly, if the 8-plex ions are determined to be present then the run is classified as 8-plex labeled rather than 4-plex labeled. Similarly, TMT comes in 2-plex, 6-plex and 10-plex varieties,^11^ and each variety’s reporter ions are a subset of the next variety. 2-plex TMT is rarely used, and when 6-plex or 10-plex TMT is used, reporter ions representing as few as three of the available channels may be present. Therefore, if any 6-plex ions that are not also 2-plex ions are determined to be present, then the experiment is labeled “6-plex or 10-plex,” rather than “2-plex.”

Although the 2-plex and 6-plex reporter ions have different nominal masses, four of the 6-plex ion nominal masses are duplicated in the 10-plex reporter ions and are only distinguishable with a high-accuracy measurement. Therefore, if 6-plex ions are detected, then a separate test is run to determine whether 10-plex ions are also present. This test consists of counting the peaks within a small *m/z* range (0.003 Da) of each reporter ion and calculating the proportion of that count over a count of peaks for each nominal mass. If, for two or more reporter ion nominal masses, the small range around the 10-plex reporter ions makes up more than 10% of the number of peaks in the window, then the run is classified as 10-plex labeled.

A further complication of TMT 10-plex labeling is that, due to the small mass difference between label masses, reporter ions are sometimes measured in MS3 spectra (e.g., in the “SPS-MS3” analysis workflow). In this case, they may not be detectable in the associated MS/MS spectra if the detector used for MS/MS is not capable of measuring the low mass range. Therefore, if MS3 spectra are present in a run, then they are evaluated for the presence of TMT 10-plex reporter ions in the same way described above for MS/MS spectra, even if no TMT reporter ions are observed in the run’s MS/MS spectra.

If iTRAQ or TMT reporter ions are detected in a run, then Param-Medic suggests searching with a variable n-terminal modification appropriate to the specific label detected.

### 2.3 Detecting SILAC Labeling

SILAC labeling is a popular method for comparing two or three samples using labels incorporated into lysine and/or arginine residues. Accordingly, in a SILAC-labeled mixed sample, the same peptide occurring in two differently-labeled samples will have a different precursor mass in each sample, potentially generating a pair of MS/MS spectra nearby in retention time, with precursor masses having a nominal mass difference of 4 Da, 6 Da, 8 Da or 10 Da, depending on the labels used.

For each SILAC mass separation, Param-Medic counts the number of such pairs separated by no more than 50 scans and calculates a Z-score comparing the observed number of pairs to the counts of pairs found at six different control mass separations (11, 14, 15, 21, 23, and 27 Da). If the Z-score exceeds an empirically-determined threshold, then the run is determined to contain the corresponding SILAC label, and Param-Medic suggests that the run be searched with an appropriate variable modification on both lysine and arginine residues.

### 2.4 Data sets

To select decision thresholds for each statistic, and to evaluate the performance of the resulting classifiers, we downloaded experiments containing each type of modification from the MassIVE proteomics repository (Table 1). We used individual filenames and information from the associated manuscripts to determine which specific runs in each experiment contained the modification and which did not. For each modification type, we assembled a list of runs not containing the modification from all of the appropriate datasets for the other modification types and the unmodified runs, if any, from the experiment containing the modification in question. In this way, we assembled a training set with a total of 196 runs and a test set with a total of 174 runs, ensuring that all runs from a given experiment are assigned either to the training or test set. Some experiments contained hundreds of runs; to ensure that no experiment dominated the test set, in those cases we chose a subset of runs that came first in lexicographic order, in an attempt to use data from similar types of samples.

**Table 1:** Experiments used in the construction and assessment modification-detection classifiers. All accession numbers are MassIVE accessions except PXD002053, which is a PRIDE accession. The score threshold determined for the 6 Da SILAC classifier, on 6 Da SILAC data, was used for all four SILAC classifiers; hence, no training data are listed for the other three. Experiment MSV000079583 had SILAC-labeled runs, iTRAQ-labeled runs and phosphorylated runs but was only used for training the iTRAQ classifier. There were no test data for 10-plex TMT, and results below are from evaluation of training data.

| Modification | Training data set(s) | Test data set(s) |
| --- | --- | --- |
| SILAC (4 Da) | None | MSV000079097<sup>12</sup> |
| SILAC (6 Da) | MSV000080791<sup>13</sup> | MSV000079097, MSV000079583<sup>14</sup> |
| SILAC (8 Da) | None | MSV000079097 |
| SILAC (10 Da) | None | MSV000079097, MSV000079583 |
| TMT (6-plex) | PXD002053<sup>15</sup> | MSV000080777,<sup>16</sup> MSV000080710<sup>17</sup> |
| TMT (10-plex) | MSV000079068<sup>17</sup> | None |
| iTRAQ (4-plex) | MSV000079583 | MSV000082884<sup>18</sup> |
| iTRAQ (8-plex) | MSV000080696<sup>19</sup> | MSV000080848<sup>20</sup> |
| Phosphorylation | MSV000080773 | MSV000079583, MSV000080777 |

### 2.5 Selecting decision thresholds

To select a decision threshold for each of the statistics described above, we ran Param-Medic on all of the training set runs. In each case, a higher value of the statistic indicates a stronger signal representing the presence of the modification. For each statistic, we selected a decision threshold that maximized classification accuracy on the training set runs.

### 2.6 Evaluation

Finally, we ran Param-Medic on the test set runs. For each classifier, we considered the runs containing the associated modification to be positive examples and the remaining runs to be negative examples. We created a receiver operator characteristic (ROC) curve describing the performance of the classifier, and we computed sensitivity and specificity at the selected decision boundary.

## 3 Results

Overall, evaluation of Param-Medic on the test set suggests that the software performs very well across the 10 binary classification tasks that we considered (Table 2 and Figure 1).

**Table 2:** Performance of the classifier on each modification type. Classifiers were evaluated on total runs. Some runs contained multiple modification types. “AUROC” is the area under the ROC curve.

| Modification | Runs with Modification | Sensitivity | Specificity | AUROC |
| --- | --- | --- | --- | --- |
| TMT (6-plex) | 53 | 0.98 | 0.98 | 0.98 |
| TMT (10-plex) | 5 | 0.80 | 0.86 | 0.75 |
| iTRAQ (4-plex) | 58 | 0.98 | 0.99 | 0.99 |
| iTRAQ (8-plex) | 30 | 0.97 | 1.00 | 1.00 |
| SILAC (4 Da) | 12 | 0.92 | 1.00 | 1.00 |
| SILAC (6 Da) | 37 | 0.57 | 0.99 | 0.96 |
| SILAC (8 Da) | 12 | 1.00 | 0.99 | 1.00 |
| SILAC (10 Da) | 37 | 0.89 | 0.99 | 0.98 |
| Phosphorylation | 17 | 0.94 | 0.98 | 0.95 |

**Figure 1:**
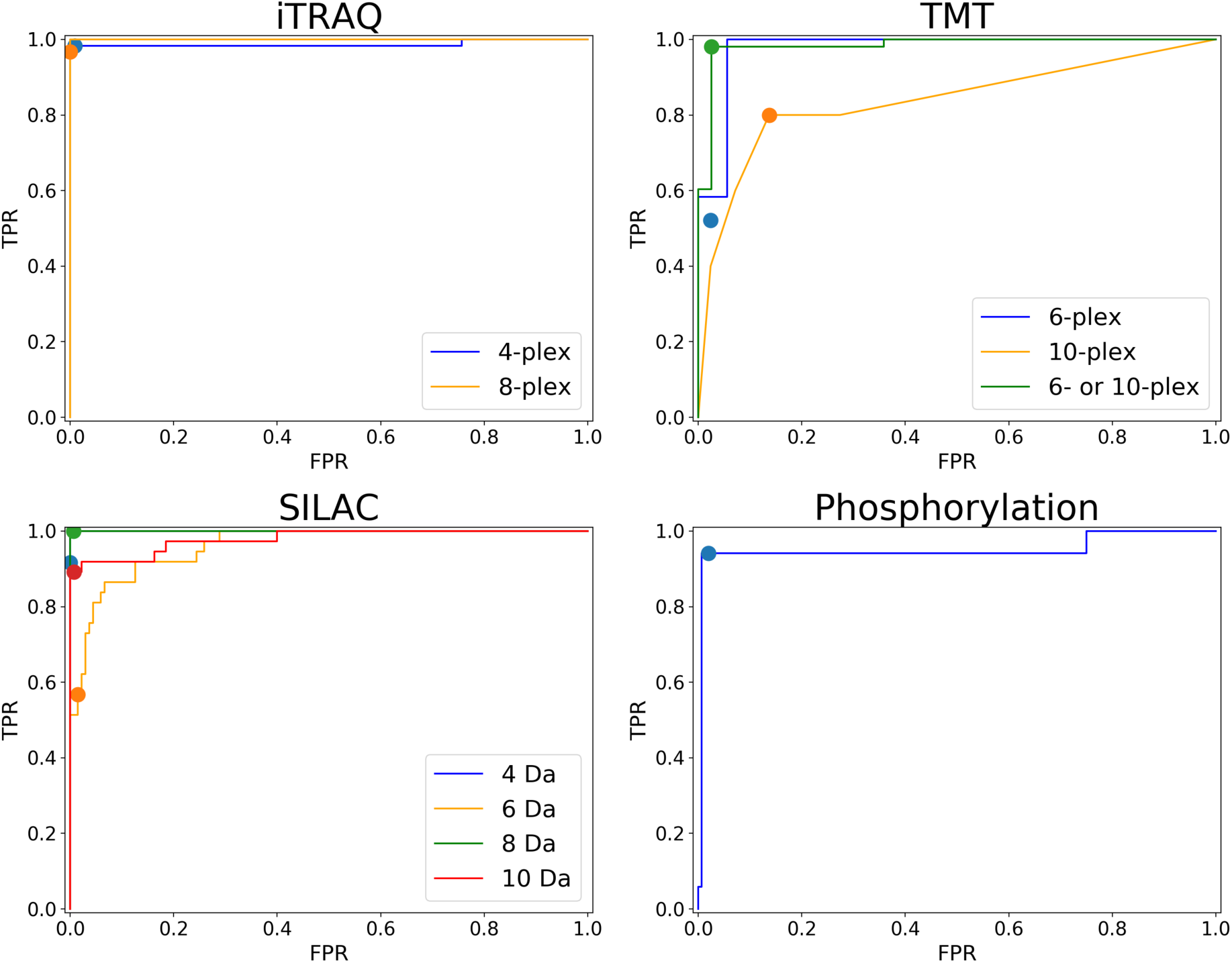
Modification detection performance. ROC curves describing the performance of each of the classifiers. The dot on each line corresponds to the selected decision threshold. All ROC curves are for test data sets, except that TMT10 is for training data.

TMT labeling can be detected accurately, except that Param-Medic has a difficult time distinguishing between 6-plex and 10-plex TMT, even on the training data. Since the two varieties of TMT require the same search parameter settings (but different quantitative analysis), detecting 6-plex TMT in run with 10-plex labeling is still valuable. Therefore, if 10-plex TMT is detected, then TMT 6-plex is also indicated as present.

In the case of iTRAQ, both settings (8-plex and 4-plex) can be identified very accurately.

For SILAC, the performance for 4 Da, 8 Da and 10 Da labeling is quite good, but classification of 6 Da SILAC labeling is not as successful, with only 57% sensitivity at 99% specificity. Looking more closely at these results, we observe a stark difference between the classifier’s performance on the 25 runs from experiment MSV000079583 (100% sensitivity), which had only 6 Da and 10 Da SILAC separations, and the 23 runs from experiment MSV000079097 (0% sensitivity), which had all four SILAC separations. The relative proportion of pairs of ions separated by 6 Da in experiment MSV000079097, although somewhat higher than background, is far weaker than the relative proportion of pairs of ions with the other three SILAC separations, and also far smaller than the relative proportion of pairs of ions separated by 6 Da in experiment MSV000079583. It may be that the other three SILAC separations interfere with the approach, or that experiment MSV000079097 simply doesn’t contain as many 6 Da-separated SILAC pairs.

Finally, we find that detection of enrichment for phosphorylated peptides is somewhat difficult, possibly due to the presence of a small proportion of phosphorylated peptides in un-enriched samples. Our classifier achieves 94% sensitivity at 98% specificity.

In the case of phosphorylation, we found ourselves using Param-Medic in precisely the way we anticipate it being used by other researchers. One of our datasets (with MassIVE accession MSV000080777) contained both TMT tags and phospho-enrichment. Initially, our phosphorylation classifier appeared to have very low sensitivity (57%) on the 28 runs from this experiment. However, upon investigation, all 11 of the files the classifier declared to be non-phosphorylated contained “_FT_” in their filenames. The associated manuscript^16^ described an analysis that may have been performed on the flow-through, or phospho-depleted portions of the samples, but the manuscript and MassIVE submission did not indicate which files represented phosphorylated samples. We therefore changed the labels of those runs in our analysis: when calculating the sensitivity and specificity that we report for the phosphorylation classifier, we assumed that “_FT_” does indeed represent the flow-through samples.

## 4 Discussion

We have described enhancements to Param-Medic that detect the presence of SILAC labeling, TMT and iTRAQ labeling, and enrichment of phosphorylated peptides with high sensitivity and specificity. These tools, combined with the existing Param-Medic estimation of precursor and fragment mass error, provide many of the parameter values needed to successfully search mass spectrometry data files whose metadata are not known *a priori*.

In fact, the classifiers described here were useful in the development of the classifiers themselves. The apparent lack of signal representing phosphorylation in 11 of the runs from one of our phosphorylated experiments, all from files with “_FT_” in the filename, led us to investigate and determine that those 11 files likely represented flow-through fractions. Param-Medic can be helpful in unraveling the puzzle of which specific runs from a publicly available experiment represent which analyses. In this way, Param-Medic could save a great deal of time for researchers engaged in large-scale meta-analyses of public data.

Param-Medic has been implemented as a standalone Python 2.7 tool which may be downloaded (including source code) at https://github.com/dhmay/param-medic or simply added to a Python installation with the “pip” tool. It has also been incorporated into version 3.2 of the Crux Toolkit, available at http://crux.ms.

## Acknowledgments

We are grateful to Nuno Bandeira for helpful discussions and for pointing us to relevant data sets in MassIVE. Research reported in this publication was supported by the National Defense Science and Engineering Graduate Fellowship (NDSEG) Program, Natioanl Science Foundation award 1633939, and National Institutes of Health award R01 GM121818.

## References

[1] May, D. H.; Tamura, K.; Noble, W. S. Journal of Proteome Research 2017, 16, acs.jproteome.7b00028.

[2] Ong, S.-E.; Blagoev, B.; Kratchmarova, I.; Kristensen, D. B.; Steen, H.; Pandey, A.; Mann, M. Molecular & cellular proteomics : MCP 2002, 1, 376–86.

[3] Wiese, S.; Reidegeld, K. A.; Meyer, H. E.; Warscheid, B. Proteomics 2007, 7, 340–350.

[4] Pottiez, G.; Wiederin, J.; Fox, H. S.; Ciborowski, P. Journal of Proteome Research 2012, 11, 3774–3781.

[5] Thompson, A.; Schäfer, J.; Kuhn, K.; Kienle, S.; Schwarz, J.; Schmidt, G.; Neumann, T.; Hamon, C. Analytical Chemistry 2003, 75, 1895–1904.

[6] Dayon, L.; Hainard, A.; Licker, V.; Turck, N.; Kuhn, K.; Hochstrasser, D. F.; Burkhard, P. R.; Sanchez, J. C. Analytical Chemistry 2008, 80, 2921–2931.

[7] Mingxun, W.; Wang, J.; Carver, J.; Pullman, B. S.; Cha, S. W.; Bandeira, N. Cell Systems 2018, 7, 412–421.

[8] Fílla, J.; Honys, D. Enrichment techniques employed in phosphoproteomics. 2012.

[9] Lehmann, W. D.; Krüger, R.; Salek, M.; Hung, C. W.; Wolschin, F.; Weckwerth, W. Journal of Proteome Research 2007, 6, 2866–73.

[10] Wolski, W. E.; Farrow, M.; Emde, A.-K.; Lehrach, H.; Lalowski, M.; Reinert, K. Proteome science 2006, 4, 18.

[11] Werner, T.; Sweetman, G.; Savitski, M. F.; Mathieson, T.; Bantscheff, M.; Savitski, M. M. Analytical Chemistry 2014, 86, 3594–3601.

[12] Udeshi, N. D.; Svinkina, T.; Mertins, P.; Kuhn, E.; Mani, D. R.; Qiao, J. W.; Carr, S. A. Molecular & Cellular Proteomics 2013,

[13] Jiang, Y.; Lee, J.; Lee, J. H.; Lee, J. W.; Kim, J. H.; Choi, W. H.; Yoo, Y. D.; Cha-Molstad, H.; Kim, B. Y.; Kwon, Y. T.; Noh, S. A.; Kim, K. P.; Lee, M. J. Autophagy 2016, 12, 2197–2212.

[14] Shiromizu, T.; Adachi, J.; Watanabe, S.; Murakami, T.; Kuga, T.; Muraoka, S.; Tomonaga, T. Journal of Proteome Research 2013, 12, 2414–21.

[15] Clark, D. J.; Fondrie, W. E.; Liao, Z.; Hanson, P. I.; Fulton, A.; Mao, L.; Yang, A. J. Analytical Chemistry 2015, 87, 10462–10469.

[16] Hoffman, N. J. et al. Cell metabolism 2015, 22, 922–35.

[17] Xu, B. et al. Neurobiology of Aging 2016, 39, 46–56.

[18] Sahasrabuddhe, N. A.; Barbhuiya, M. A.; Bhunia, S.; Subbannayya, T.; Gowda, H.; Advani, J.; Shrivastav, B. R.; Navani, S.; Leal, P.; Roa, J. C.; Chaerkady, R.; Gupta, S.; Chatterjee, A.; Pandey, A.; Tiwari, P. K. Biochemical and Biophysical Research Communications 2014, 446, 863–9.

[19] Guo, J.; Ren, Y.; Hou, G.; Wen, B.; Xian, F.; Chen, Z.; Cui, P.; Xie, Y.; Zi, J.; Lin, L.; Wu, S.; Li, Z.; Wu, L.; Lou, X.; Liu, S. Journal of Proteome Research 2016, 15, 2164–77.

[20] Preil, S. A.; Kristensen, L. P.; Beck, H. C.; Jensen, P. S.; Nielsen, P. S.; Steiniche, T.; Bjørling-Poulsen, M.; Larsen, M. R.; Hansen, M. L.; Rasmussen, L. M. Circulation: Cardiovascular Genetics 2015, 8, 727–35.

